# Connectivity profile and function of uniquely human cortical areas

**DOI:** 10.1101/2024.06.20.599486

**Authors:** Katherine L. Bryant, Julia Camilleri, Shaun Warrington, Guilherme Blazquez Freches, Stamatios N. Sotiropoulos, Saad Jbabdi, Simon Eickhoff, Rogier B. Mars

## Abstract

Quantitative comparison of the white matter organization of the human neocortex with that of the chimpanzee and macaque shows a wide distribution of areas with a uniquely human connectivity profile, including the frontal-parietal fiber systems and the temporal visual pathway. Functional decoding of these areas shows their involvement in language, abstract reasoning, and social information processing. Overall, these results counter models that assign primacy to prefrontal cortex for human uniqueness.

## Main Text

Our human behavioral repertoire enables us to spread across the globe into a much greater variety of niches than any other primate. Various behavioral innovations have alternatively been suggested to characterize our abilities, including our collaborative social abilities, tool use, ability for mental time travel, and spoken language^1–3^. Understanding the basis of uniquely human behavior requires a comparison of our brain to that of our closest primate relatives. Such comparisons tend to focus on measures of size, highlight that the human neocortex or cerebellum is expanded^4^, that certain areas are preferentially expanded^5^, or that the absolute number of neurons in the human brain outstrips that of other primates^6^. None of these measures, however, provides a link to the behavior that the brain produces and that, ultimately, is the likely target of selection. In contrast, work in neuroimaging has highlighted measures of brain organization at the level of areal connections that do have predictive value regarding the function of parts of the brain^7,8^. Hence, the level of large-scale connections between brain areas is a more suitable level of between-species comparison of brain organization if one wants to understand the unique abilities of the human brain in the context of other primates.

Connectivity can now be studied at the whole brain level using diffusion MRI and associated tractography algorithms, offering a new type of data for comparative and evolutionary neuroscience^9^. Recent work has created standardized protocols for reconstructing the major fiber pathways of the primate brain, creating white matter atlases of the human, the developing human, and the macaque monkey brain^10,11^. These methods characterize the cortical areas of each species’ brain in terms of its connectivity with major white matter bundles, known to be homologous among primates. By describing all cortical areas of all brains in terms of connectivity to homologous tracts, we, in effect, place all the brains within a *common connectivity space*. This allows a quantitative comparison of brain organization across species^12^. While previous studies focused on comparisons of the human brain with that of the most-often studied primate, the macaque, here we exploit our recently developed comparable comprehensive white matter atlases of the chimpanzee^13^ allowing us to directly compare humans with our closest relatives, as well as the macaque.

We describe each point on the cortical surface of the human and chimpanzee brain as a vector of connectivity probabilities with 18 white matter fiber bundles that are homologous across species. Given that the connectivity profiles are anchored on homologous white matter fibers, the connectivity pattern of a human vertex can then be compared to that of each chimpanzee and macaque vertex by calculating the Kullback-Leibler (KL) divergence between connectivity profiles^10^. The best matching vertex in the non-human species is the one with the minimum KL value. Overall spatial maps of divergence of the human brain to the chimpanzee is then visualised by plotting the minimum KL value for each human vertex. This shows large zones of divergence in the middle temporal lobe, temporoparietal cortex, and lateral frontal cortex with a particular hotspot in the dorsal frontal cortex (Fig. 1, left).

**Figure 1.**
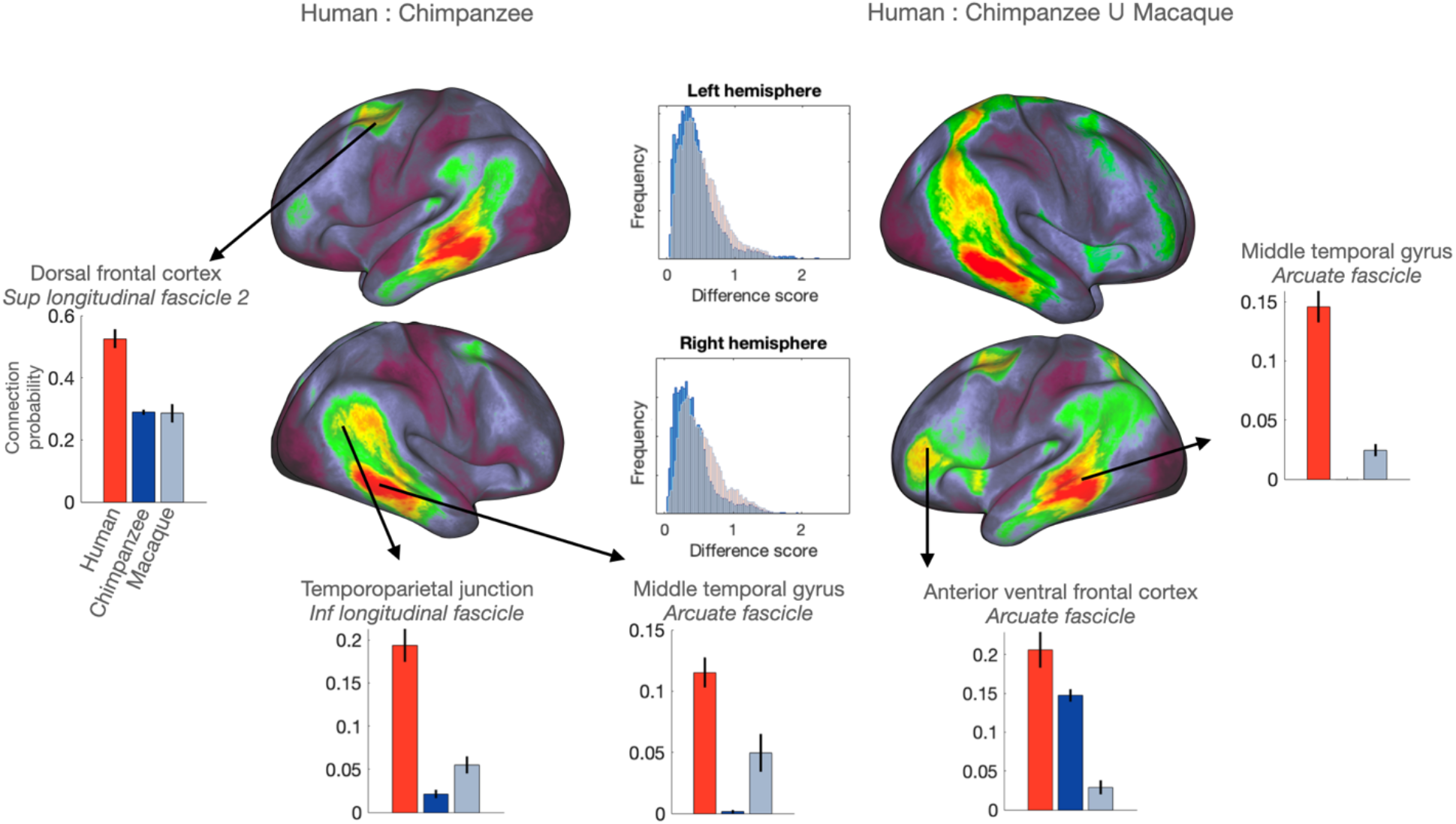
Mapping connectivity divergence between primates identifies multiple hotspots of human specialization. Divergence maps of the human brain showing vertices with connectivity profiles that have a poor match in the chimpanzee (left) in either the chimpanzee or the macaque. Bar graphs show the normalized connectivity (±SEM) of the selected vertex with a tract driving these differences in the human (red) and of its best matching vertices in the chimpanzee (dark blue) and macaque (light blue). Tracts include SLF2 (superior longitudinal fascicle 2), ILF (inferior longitudinal fascicle) and AF (arcuate fascicle). Full connectivity profiles of the human vertices and their best matches in the other species in the Supplementary Material. Histograms in the center show the distribution of KL values comparing human and chimpanzee (blue) and human and macaque (red).

The divergence of the human brain from the chimpanzee brain can be compared to the divergence of the human brain with the macaque. The distribution of minimum KL values when comparing the human and the chimpanzee differs from that when comparing the human and the macaque (Kolmogorov-Smirnov (K-S) test *p*<0.001 for both hemispheres). Plotting the distribution of minimum KL values based on the union of KL values with the chimpanzee and the macaque indeed shows broader differences, with increases in divergence in the anterior ventral frontal cortex and posterior parietal cortex (Fig 1, middle).

Having established *which* cortical areas show the greatest divergence between species, we could then assess *how* the connectivity profile of areas of human divergence differs from that of the closest match in the other species by identifying which connections are driving the observed differences in organization. Further, we can use meta-analytic data of functional brain activation to investigate the functional *roles* of divergent regions in the human brain, linking anatomical differences between species’ brains to behavior.

Divergence between the human brain and both the chimpanzee and macaque were evident in the dorsal frontal cortex. The vertices of high divergence overlap with anterior area 6, the inferior 6-8 transition area, and the frontal eye fields^14^. The connectivity profile of this area is dominated by the frontal-parietal superior longitudinal fascicle, in particular the second branch (SLF2)^15^ (Fig. 1; see Suppl. Fig. 1 for full connectivity profiles). Using the common connectivity space, we can determine which vertices in the chimpanzee and the macaque have a connectivity profile that is the least different from that of the human. Extracting the connectivity of these vertices shows that even these do not show strong SLF2 connectivity (Fig. 1, Suppl. Fig. 2). We thus conclude that strong SLF2 connectivity in this part of dorsal frontal cortex is driving the divergence in brain organization between the human and the other two primates.

To assess the functional role of these regions, we turned to a database of functional neuroimaging studies (brainmap.org)^16^. We assessed if, for a given behavioral domain, the probability of finding activation of a region was significantly higher than the *a priori* chance, so-called *forward inference*. This approach allows a functional characterization of the areas we identified as structurally divergent from other primate brains (Fig. 2, see Supplemental Material for full decoding results). For the three dorsal frontal regions mentioned above, the behavioral domains most likely to activate them include spatial cognition, working memory, and reasoning. Some of these regions have previously been identified as part of the so-called multiple demand network^17^, a network of mostly parietal and frontal regions that consistently activate for a range of high-level cognitive tasks. Although homologs of this network exist in the macaque, recent comparative work shows that the connections between these regions are much more extensive in the human^18^. As such, it has been suggested that human domain-general knowledge has a precursor in parietal-frontal network originally evolved for visuomotor control in early primates^19^. The current results extend this finding to our nearest animal relative, and directly link anatomical differences to functional domains associated with the multiple demand network.

**Figure 2.**
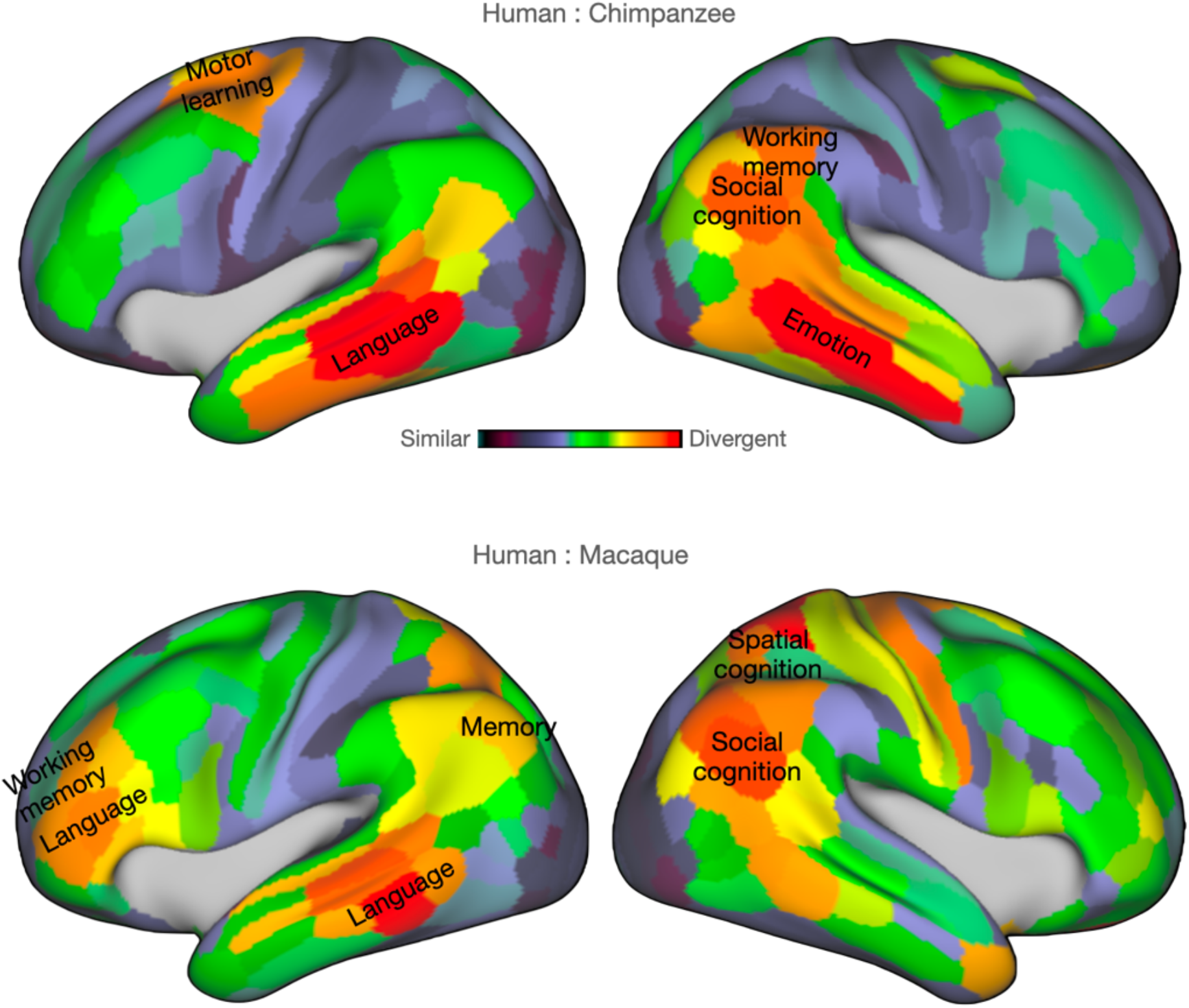
Decoding areas of high divergence highlights multiple behavioural domains. Functional activations that correlate most with areas of high KL divergence for the human : chimpanzee comparison (A) and the human : macaque comparison (B). Areas of high divergence are parcellated according to Glasser and colleagues^14^.

Extensive differences between the human and non-human brains were found in ventral frontal cortex and middle temporal gyrus. Both these hotspots of divergence were driven by more extensive connectivity of the arcuate fascicle (AF) in humans (Fig. 1). Such AF connectivity in the human brain has been shown before^20,21^, but the comparison of the human with the chimpanzee on the one hand and the chimpanzee and macaque on the other shows a dissociation between frontal and temporal cortex. While the best matching vertices for middle temporal cortex showed a lack of innervation of the AF in both chimpanzees and macaques, the best matching vertices to the anterior ventral frontal cortex show some AF in the chimpanzee, but none in the macaque. This suggests a scenario where the extension of the AF occurred gradually, with frontal expansions occurring in the ape lineage, preceding temporal expansions into the middle temporal cortex in the human lineage.

Consistent with the role of the AF in human language, functional decoding of both the middle temporal and ventral frontal cortex in the left hemisphere yielded the behavioral domain ‘language’ prominently. However, it was clear that the AF extension, especially in the temporal cortex, was bilateral. Decoding of the right middle temporal cortex yielded the domain ‘emotion’. Although the function of right temporal association cortices is yet not well-characterized in the fMRI literature, lesion studies suggest they play a role in nonverbal semantic social cognition^22^. Importantly, these results speak against a language-only interpretation of AF extensions in the ape and human brain.

A prominent zone of divergence between the human brain and that of both the chimpanzee and macaque was in the posterior superior temporal cortex and inferior parietal lobule, together often referred to as the temporoparietal junction area (TPJ). The posterior TPJ especially has often been associated with the human ability to entertain others’ belief states, so-called mentalizing or Theory of Mind^23^. The hotspot of divergence overlaps with this area, and functional decoding indeed shows ‘social cognition’ as its most significant behavioral domain. The human posterior TPJ shows strong connectivity to the inferior longitudinal fascicle (ILF), which is not present in the other two species (suppl. Fig. 6). The ILF is part of the ventral visual pathway but extends into parietal cortex in anthropoid primates^24^. It is thought that the ILF has expanded in great apes and that the dorsal component has a role in social cognition, allowing some of the temporal cortex machinery for visual processing to be adapted for social information processing^25,26^. The current results connect these two findings of TPJ’s role in social cognition and ILF’s prominent expansion by showing the TPJ is innervated by the ILF in the human.

Our results thus argue against a single explanatory factor or evolutionary event driving the uniquely human behavioral repertoire. While current theories on human brain uniqueness focus on changes to prefrontal areas, our findings support a two-step evolutionary process, in which changes in prefrontal cortex organization emerge prior to changes in temporal areas. Unlike global connectivity or gross anatomical approaches, anatomically-informed comparative connectivity makes it possible to reveal major changes in multiple association fiber systems underlying a variety of cognitive functions that have changed in a stepwise manner in the great ape and human lineages.

## Acknowledgements

The work of RBM and KLB was supported by the Biotechnology and Biological Sciences Research Council UK [BB/N019814/1] to RBM. RBM was supported by the EPA Cephalosporin Fund. The Wellcome Centre for Integrative Neuroimaging is supported by core funding from the Wellcome Trust [203129/Z/16/Z]. Chimpanzee brain scans were acquired prior to the 2015 implementation of U.S. Fish and Wildlife Service and National Institutes of Health regulations governing research with chimpanzees and made possible through the Yerkes Base Grant [ORIP/OD P51OD011132]. For the purpose of Open Access, the author has applied a CC BY public copyright licence to any Author Accepted Manuscript version arising from this submission.

## References

1. Healy, S.D. (Oxford University Press: Oxford, 2021).

2. Suddendorf, T., Redshaw, J. & Bulley, A. (Basic Books: New York, 2022).

3. Tomasello, M. & Vaish, A. Annu Rev Psychol 64, 231–255 (2013).

4. Barton, R.A. & Venditti, C. Curr Biol 24, 2440–2444 (2014).

5. Donahue, C.J., Glasser, M.F., Preuss, T.M., Rilling, J.K. & Van Essen, D.C. Proc Natl Acad Sci USA 115, E5183–E5192 (2018).

6. Herculano-Houzel, S. PNAS 109, 10661–10668 (2012).

7. Mars, R.B., Passingham, R.E. & Jbabdi, S. Trends Cogn Sci 22, 1026–1037 (2018).

8. Saygin, Z.M. et al. Nat Neurosci 19, 1250–1255 (2016).

9. Thiebaut de Schotten, M. & Forkel, S.J. Science 378, 505–510 (2022).

10. Mars, R.B. et al. eLife 7, e35237 (2018).

11. Warrington, S. et al. Sci Adv 8, eabq2022 (2022).

12. Mars, R.B., Jbabdi, S. & Rushworth, M.F.S. Annu Rev Neurosci 44, 69–86 (2021).

13. Bryant, K.L., Li, L., Eichert, N. & Mars, R.B. PLoS Biol 18, e3000971 (2020).

14. Glasser, M.F. et al. Nature 536, 171–178 (2016).

15. Thiebaut de Schotten, M. et al. Nat Neurosci 14, 1245–1246 (2011).

16. Fox, P.T. et al. Hum Brain Mapp 25, 185–198 (2005).

17. Assem, M., Glasser, M.F., Van Essen, D.C. & Duncan, J. Cereb Cortex 30, 4361–4380 (2020).

18. Karadachka, K. et al. Cereb Cortex 33, 10959–10971 (2023).

19. Genovesio, A., Wise, S.P. & Passingham, R.E. Trends Cogn Sci 18, 72–81 (2014).

20. Rilling, J.K. et al. Nat Neurosci 11, 426–428 (2008).

21. Sierpowska, J. et al. Proc Natl Acad Sci U S A 119, e2118295119 (2022).

22. Borghesani, V., et al. Cortex, 115, 2-85 (2019).

23. Schurz, M., Tholen, M.G., Perner, J., Mars, R.B. & Sallet, J. Hum Brain Mapp 38, 4788–4805 (2017).

24. Roumazeilles, L. et al. Cereb Cortex 32, 1608–1624 (2022).

25. Pitcher, D. & Ungerleider, L.G. Trends Cogn Sci 0, (2020).

26. Roumazeilles, L. et al. PLoS Biol 18, e3000810 (2020).

