## Supplemental Materials for "Connectivity profile and function of uniquely human cortical areas"

### Data

#### *Human*

Thirty human subjects (16 female, ages 22-35) were selected from the *in vivo* diffusion MRI data provided by the Human Connectome Project (HCP), WU-Minn Consortium (Principal Investigators: David Van Essen and Kamil Ugurbil; 1U54MH091657) funded by 16 NIH Institutes and Centers and the McDonnell Center for Systems Neuroscience at Washington University[1]. Minimally preprocessed datasets from the Q2 public data release were used. Data acquisition and preprocessing methods have been previously described[2,3]. Briefly, 1.25 mm isotropic resolution diffusion-weighted data were collected on a 3T Siemens Skyra scanner with a slice-accelerated gradient echo EPI readout. *Q*-space sampling included 3 shells at $b$ = 1000, 2000, and 3000 s/mm$^2$. Ninety diffusion encoding gradient directions and 6 b = 0's were obtained twice for each shell, with the phase-encoding direction reversed. An MPRAGE sequence was used to acquire T1-weighted (T1w) images at .7 mm isotropic resolution, then aligned to diffusion space using the HCP minimal preprocessing pipeline[2]. Diffusion-weighted images were processed with FSL, using FMRIB's Diffusion Toolbox and bedpostX[4,10]. A high-resolution 164k surface mesh (~164,000 vertices per hemisphere), as well as a lower-resolution mesh (32k) were generated using the PostFreeSurfer pipeline.

#### *Chimpanzee*

Chimpanzee (*Pan troglodytes*; n=23, 26 ± 11 yrs, all female) MR scans were obtained from an archive hosted by the National Chimpanzee Brain Resource. Scans acquired prior to the 2015 implementation of U.S. Fish and Wildlife Service and National Institutes of Health regulations governing research with chimpanzees. All the scans reported here were collected as part of a grant to study aging in female primates, were completed by 2012, and have been used in previous studies[5,6]. Chimpanzees were housed at the Emory National Primate Research Center (ENPRC, Atlanta, Georgia, USA) and all procedures were carried out in accordance with protocols approved by the ENPRC and the Emory University Institutional Animal Care and Use Committee (IACUC approval #YER-2001206).

Following standard ENPRC veterinary procedures, chimpanzee subjects were immobilized with ketamine injections (2–6 mg/kg, i.m.), then anesthetized with an intravenous propofol drip (10 mg/kg/h) prior to scanning. Subjects remained sedated for the duration of the scans as well as the time required for transport between the scanner and their home cage. Primates were housed in a single cage for 6–12 h after scanning to recover from the effects of anesthesia before being returned to their home cage and cage mates. Veterinary and research staff evaluated the well-being of chimpanzees twice daily after the scan for possible post-anesthesia distress.

MR scanning protocols and preprocessing for the chimpanzee dataset have been described in detail previously[7,8] Briefly, anatomical and diffusion MR scans were acquired *in vivo* in a Siemens 3T Trio scanner (Siemens Medical System, Malvern, PA, USA). Diffusion-weighted MRI data were collected with a single-shot, pulsed-gradient spin-echo echo-planar imaging sequence. Parameters were as follows: 41 slices were scanned at a voxel size of 1.8 $mm^3$, TR/TE: 5900 ms/86 ms, matrix size: 72x128. Two diffusion-weighted images were acquired for each of 60 diffusion directions (b=1000 s/mm2), each with 1 of the possible left–right phase-encoding directions and 8 averages, allowing for correction of susceptibility-related distortion[9]. For each averaged diffusion-weighted image, six images without diffusion weighting (b = 0 s/$mm^2$) were also acquired. High-resolution T1w images were acquired with a MPRAGE sequence. Diffusion-weighted images were processed using FMRIB's Diffusion Toolbox and bedpostX[4,10]. Template generation for chimpanzees (previously described in detail[11,12]); involved the PreFreeSurfer pipeline was used to align the T1w and T2w volumes of 29 individual chimpanzees to native anterior commissure-posterior commissure space. Cortical surfaces and registrations to a population-specific chimpanzee template were generated using a modified version of the HCP minimal preprocessing pipeline[2]. The PostFreeSurfer pipeline was used to produce a high-resolution 164k surface mesh and a lower-resolution mesh (20k).

*Macaque*

Eight post mortem macaque brain scans (*Macaca mulatta*, n=8; six male, age range 4-14 years) were acquired using a 7T magnet with an Agilent DirectDrive console (Agilent Technologies, Santa Clara, CA, USA). Acquisition and preprocessing have been detailed previously[8]. In brief, a 2D diffusion-weighted spin-echo protocol was implemented (DW-SEMS, TE/TR: 25 ms/10 s; matrix size: 128 x 128; resolution: 0.6 x 0.6 mm; number of slices: 128; slice thickness: 0.6 mm). Nine non-diffusion-weighted (b=0 s/$mm^2$) and 131 diffusion-weighted (b=4000 s/$mm^2$) volumes were acquired with diffusion directions distributed over the whole sphere. The b=0 images were averaged and spatial signal inhomogeneities were restored. *Ex vivo* tissue usually has reduced diffusivity, necessitating larger b-values to achieve equivalent diffusion contrast to *in vivo* data; this was achieved here by increasing the diffusion sensitization from b = 1000 to 4000 s/$mm^2$. Diffusion-weighted images were processed using the same method as chimpanzees, described above. The cortical surface of one macaque with high quality structural MRI was reconstructed using a modified version of the HCP pipeline, nonlinearly registered to the other brains using FSL's FNIRT, warped to the other macaque brains and transformed to F99 standard space[13].

### Between-species comparison based on white matter tracts

*Reconstruction of major white matter tracts*

Eighteen major white matter bundles were reconstructed for all three species using probabilistic tractography[10]. A set of standardized masks previously developed for the human, chimpanzee, and macaque[8] were used to reconstruct tracts based on objective anatomical landmarks that could be identified in all species. The logic behind this approach is that a set of seed, waypoint, stop, and exclusion masks are used to define the body of any white matter tract; the tractography algorithm is then free to reconstruct the rest of the bundle, including its grey matter termination points. In this way, we have something we can objectively define as homologous across the species (the body of the tract based on anatomical criteria) and something that varies across species and is the target of our investigation (the grey matter terminations).

All combinations of seed, waypoint, stop, and exclusion masks are described in detail in previous communications. The white matter tracts studied in the present study were the anterior commissure (AC), arcuate fascicle (AF), peri-genual, dorsal, and temporal subdivisions of the cingulum bundle (CBP, CBD, and CBT), corticospinal tract (CST), frontal aslant (FA), forceps major (FMA), forceps minor (FMI), fornix (FX), inferior fronto-occipital fascicle (IFO), inferior longitudinal fascicle (ILF), middle longitudinal fascicle (MdLF), first, second, and third branches of the superior longitudinal fascicle (SLF1, SLF2, and SLF3), uncinate fascicle (UNC), and vertical occipital fascicle (VOF).

*Common connectivity spaces and between-species comparison*

To assess the connectivity of each part of the cortical surface with each white matter fiber bundle we created (tract) x (surface) matrices which we term *common connectivity spaces*. First, tractography is performed from the cortical surface towards the whole brain white matter, creating a (surface) x (brain) matrix of connectivity. Then, each tract's tractogram is multiplied by the (surface) x (brain) matrix, resulting in the (tract) x (surface) connectivity blueprint. The rows of this blueprint represent the surface projection of each tract and the column of the blueprint present the connectivity profile of each vertex of the cortical surface. This method was first applied by Mars and colleagues[14] and is now implemented in FSL's XTRACT tool[15].

Blueprints were averaged across subjects in each species to create a species-specific connectivity blueprint. Connectivity profiles can be compared across species by calculating the (vertex) x (vertex) KL divergence between two common connectivity spaces. The best match of a vertex in one species is then found by finding the vertices with the lowest KL value (<2) in the other species. A spatial map of divergence of connectivity of one brain compared to another can be established by assigning to each vertex of the first brain the smallest KL value (minKL) across all vertices in the second brain.

### Functional Decoding

To assess the functional roles of the areas of the human cortex that showed the greatest difference with the chimpanzee and the human, we used BrainMap; a publicly available meta-analytic database of functional activation studies (www.brainmap.org[16]). BrainMap uses a structured standardized coding scheme to describe published human functional neuroimaging results. In particular, Behavioral Domains are categories and subcategories that aim to classify the cognitive functions likely to be isolated by any experimental contrast.

Functional decoding was done as follows. First, the cortex was divided into distinct regions according to the Glasser parcellation[17]. Each region was assigned the maximum within-region divergence score, i.e. the divergence value from the vertex that had the highest minKL value in the region. Second, we queried the BrainMap database in 2019 to assign the functional profile of these regions using forward inference[18]. Using forward inference, a cluster's functional profile is determined by identifying taxonomic labels for which the probability of finding activation in the respective cluster was significantly higher than the a priori chance (across the entire database) of finding activation in that particular cluster. Significance was established using a binomial test ($p < 0.05$, FDR corrected[19]). In other words, we tested whether the conditional probability of activation given a particular label [P(Activation|Task)] was higher than the baseline probability of activating the brain region in question per se [P(Activation)].

It is important to point out that the specificity of the decoding results can only be as good as the taxonomy of the BrainMap database. Thus, our results should not be taken such that any Behavioral Domain associated with an area constitutes the unique role of that area. Rather, the Behavioral Domain indicates the involvement of the area, but does not claim the brain region is limited to that Domain.

Below, we provide two tables showing the functional decoding of regions based on high divergence between the human and the chimpanzee (Suppl. Table 1) and between the human and the macaque monkey (Suppl. Table 2). Behavioral domains for significant decoding and likelihood ratios are reported. Regions are labeled according to the atlas of Glasser and co-workers[17].

**Comparison of connectivity profiles across species based on *a priori* homologs**

The main manuscript reports the connectivity profile of regions in the human brain that show a high divergence with the chimpanzee and macaque, as well as the connectivity profile of the best matching chimpanzee and macaque brain areas (Fig. 1 and Suppl. Fig. 1). It is important to note that this analysis selects those vertices in the chimpanzee and macaque brain that have the least divergent connectivity profile, independent of their location. This allows an unbiased assessment of divergence across the different species brains. As has been shown previously, this analysis is capable of identifying homologous regions that are known to have similar connectivity profiles across species[14], but does not rely on priors and is therefore more principled than comparing known homologs across species. For completion, however, we also present comparisons of the connectivity profiles of human areas with those of known homologs in the chimpanzee and macaque. We do this for all the areas presented in Figure 1 of the main manuscript.

The left dorsal prefrontal region overlaps with anterior area 6, the inferior 6-8 transition area, and the frontal eye fields[17]. In humans, this area has much stronger connectivity to SLF2, compared to its best matching chimpanzee and macaque counterparts. We extracted the connectivity profiles of area FB in the chimpanzee[20], which has been suggested to contain the frontal eye fields[21], and macaque FEF[22]. As with the best matching vertices, the human has much stronger SLF2 connectivity than the other species (Suppl. Fig. 2).

Anterior ventral frontal cortex in the human received innervations of the arcuate fascicle (AF), which was evident to a lesser extent in the chimpanzee and absent in the macaque. The human area of maximum divergence overlaps with area IFSa of Glasser and colleagues[17]. and area IFS of Neubert and co-workers[23]. The homolog of this area in the chimpanzee is difficult to establish. We extracted the connectivity profile of a vertex in area FCBm[20] in the chimpanzee and on the posterior bank of the inferior branch of the arcuate sulcus in the macaque. In both cases, these locations are, if anything, quite posterior, and therefore more likely to detect AF connectivity than human IFS. Nevertheless, the pattern of most AF connectivity in the human, less in the chimpanzee, and very little in the macaque was replicated (Suppl. Fig. 3).

The human middle temporal gyrus shows strong AF connectivity, which is much lower even in the best matching areas in the other two species. When extracting the connectivity profile of middle temporal gyrus in the chimpanzee and macaque, this pattern of relatively reduced AF in the non-human primates is even stronger (Suppl. Figs. 4 and 5).

The right temporoparietal junction (TPJ) area in the human brain shows strong innervation of the ILF, which is not seen in the best matching vertices in the chimpanzee and macaque. The homolog of TPJ is difficult to establish. Although the area overlaps with area PGi of Glasser et al. [17], it is uncertain whether it is homologous to area PG in the macaque[24]. Mars et al.[25] identified two subregions of TPJ, which they labeled TPJp and TPJa, the posterior of which shows strong activation in social cognition tasks, similar to the decoding analysis presented in the main paper[26]. Connectivity profiles of regions in the macaque inferior parietal lobule do not show a prominent ILF, but rather the IFO and MdLF. In addition, the small macaque inferior parietal lobule shows strong connectivity with the AF, which does not extend ventrally as it does in the human, as discussed above (Suppl. Fig. 6).

**Data availability:** Analysis code and tractography recipes are available online at locations linked from the lab's website (www.neuroecologylab.org). Analysis code is also part of the MR Comparative Anatomy Toolbox (Mr Cat). Raw data are for the chimpanzee are available from the National Chimpanzee Brain Resource (www.chimpanzeebrain.org), for the macaque, they are available from the PRIME-DE repository (http://fcon_1000.projects.nitrc.org/indi/indiPRIME.html)[27], and from the Human Connectome Project (www.humanconnectome.org)[1] for the human.

**Acknowledgments:** The authors would like to thank Shaun Warrington, Saad Jbabdi, and Stam Sotiropolous for their work on XTRACT toolbox and library (fsl.fmrib.ox.ac.uk/fsl/fslwiki/XTRACT) which helped enable this project. The authors would like to thank Guilherme Freches with his assistance on generating the chimpanzee whole-brain connectivity matrix, Nicole Eichert for her assistance with the production of distance-corrected surface matrices, and Longchuan Li for his assistance with chimpanzee template generation. **Funding:** K.L.B. received funding from a Marie Skłodowska-Curie Individual Fellowship Grant MSCA-IF 750026 and was supported by the Biotechnology & Sciences Research Council (BBSRC) UK [BB/N019814/1]. R.B.M. was supported by the Biotechnology and Biological Sciences Research Council UK [BB/N019814/1] and the Netherlands Organization for Scientific Research [452-13-015]. SW and SNS are supported by an ERC Consolidator Grant [ERC CoG 101000969]. The Wellcome Centre for Integrative Neuroimaging is supported by core funding from the Wellcome Trust [203129/Z/16/Z]. **Author contributions:** **Competing interests:** Authors declare no competing interests. **Data and materials availability:** Chimpanzee brain scans were acquired prior to the 2015 implementation of U. S. Fish and Wildlife Service and National Institutes of Health regulations governing research with chimpanzees, made possible through the Yerkes Base Grant [ORIP/OD P51OD011132]; an archive of these scans has been made available through the National Chimpanzee Brain Resource [NIH NINDS NS092988]. Human brain scans were made available through the Human Connectome Project [1U54MH091657].

### Supplemental Figures

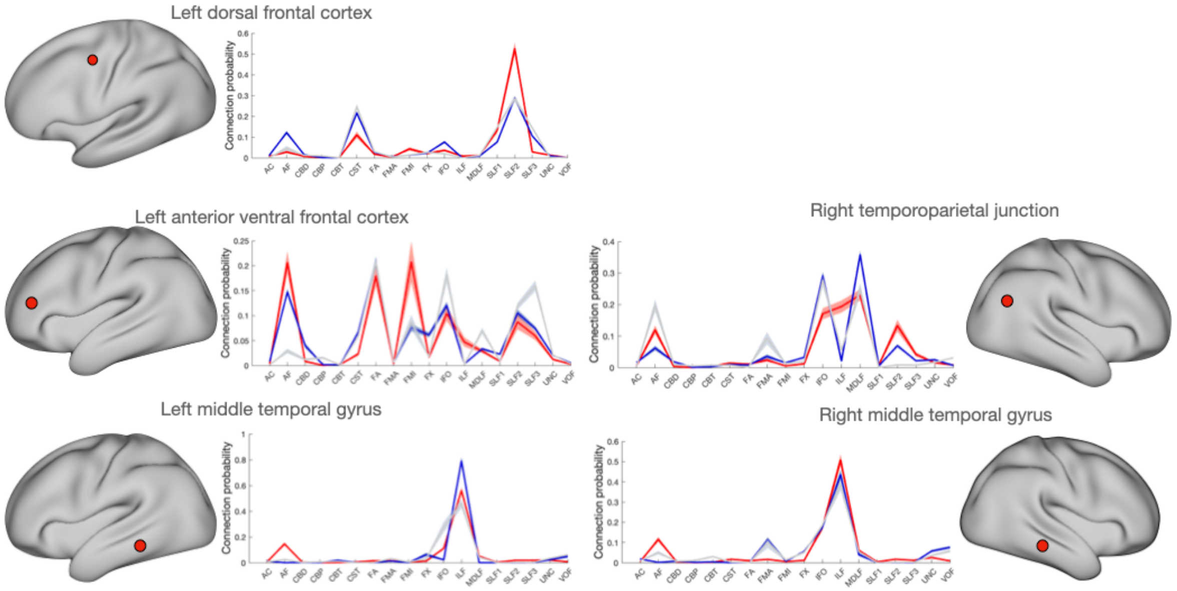


Supplementary Figure 1. Connectivity profile of areas of high between-species divergence in the human (red) and their best matching vertices in the chimpanzee (dark blue) and macaque (light blue).

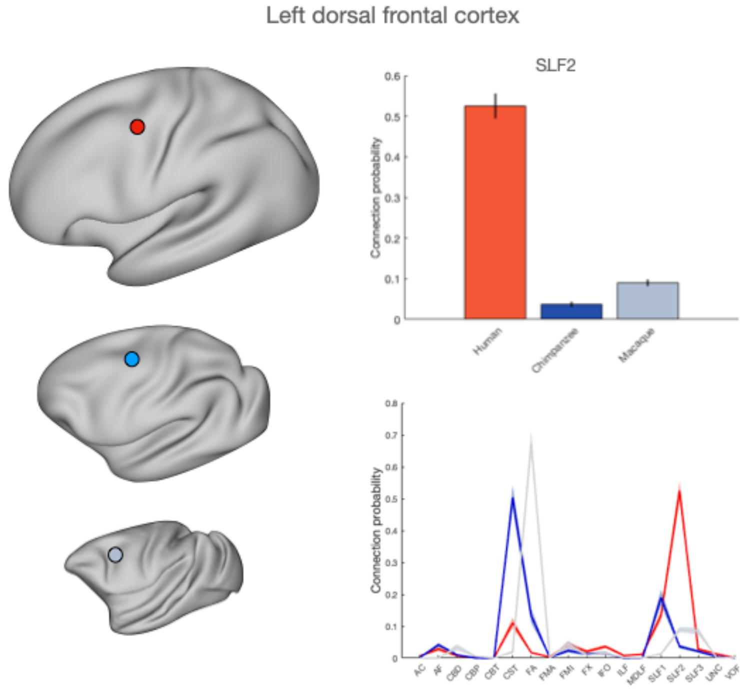


Supplementary figure 2. Connectivity of human left dorsal frontal cortex with SLF2 (top right) and with all tracts (bottom right) in red and its chimpanzee and macaque homologs in dark and light blue, respectively.

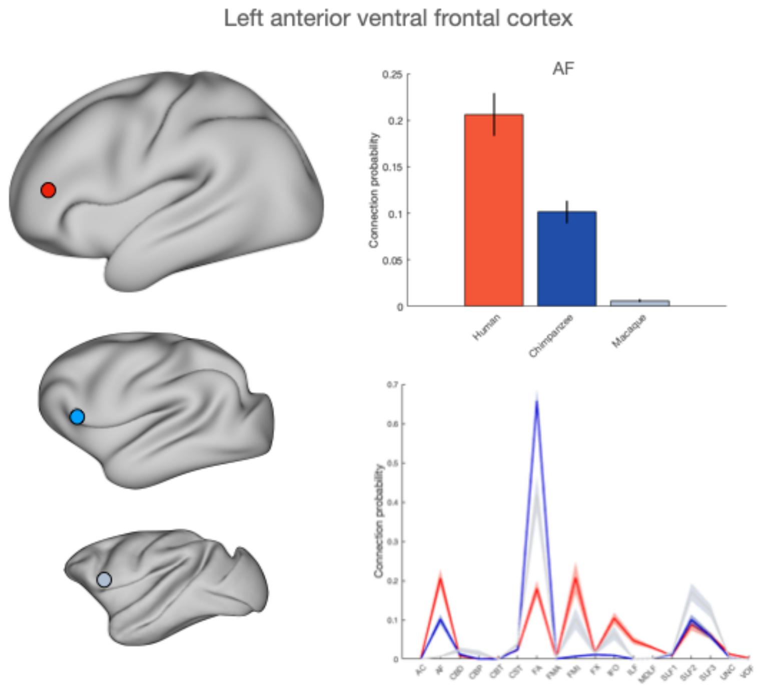


Supplementary figure 3. Connectivity of human left anterior ventral frontal cortex with AF (top right) and with all tracts (bottom right) in red and its chimpanzee and macaque homologs in dark and light blue, respectively.

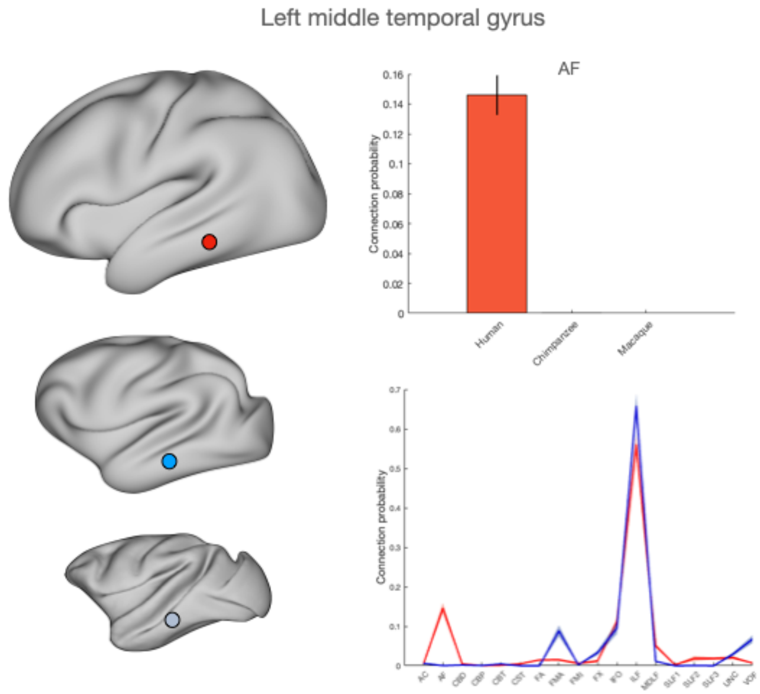


Supplementary figure 4. Connectivity of human left middle temporal gyrus with AF (top right) and with all tracts (bottom right) in red and its chimpanzee and macaque homologs in dark and light blue, respectively.

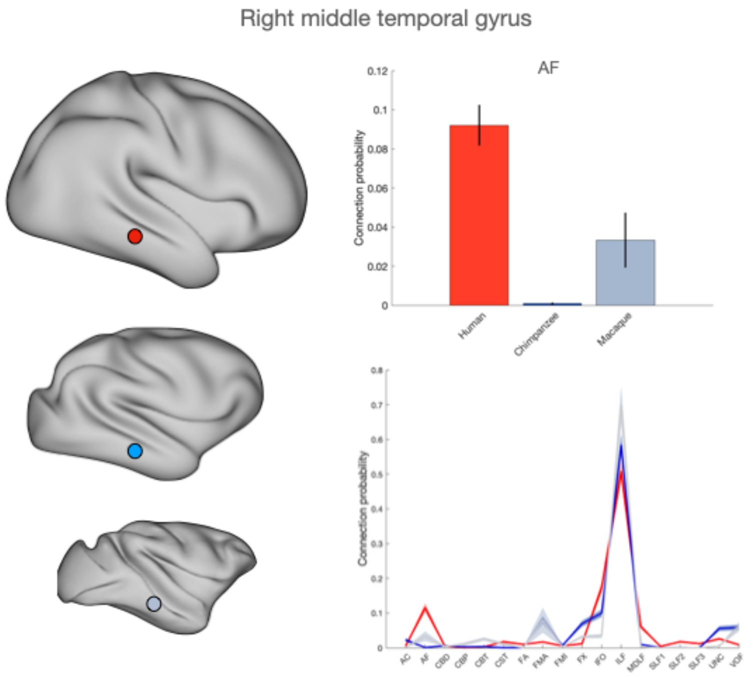


Supplementary figure 5. Connectivity of human right middle temporal gyrus with AF (top right) and with all tracts (bottom right) in red and its chimpanzee and macaque homologs in dark and light blue, respectively.

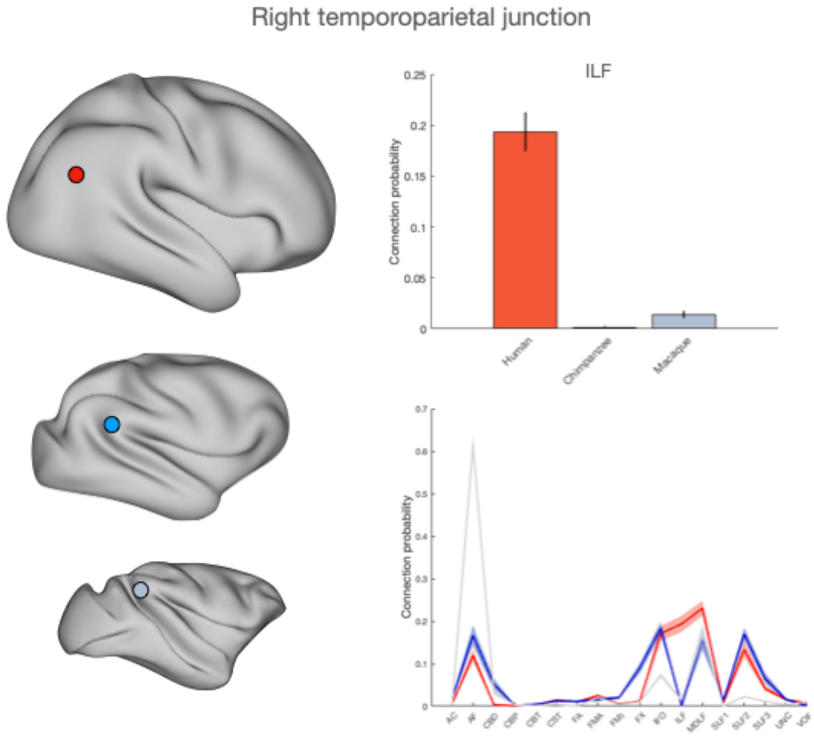


Supplementary figure 6. Connectivity of human right temporoparietal junction (TPJ) areas with ILF (top right) and with all tracts (bottom right) in red and its chimpanzee and macaque homologs in dark and light blue respectively.

### Supplementary Tables

**HUMAN : CHIMPANZEE**

**LEFT HEMISPHERE**

| *lobe* | *area* | *domain* | *LR* |
|---|---|---|---|
| **Frontal** | 6ma | Action.Imagination | 3,41 |
| | i6-8 | No significant effects | |
| | 6a | Cognition.Reasoning | 1,4 |
| | | Cognition.Memory.Working | 1,77 |
| | | Action.Execution | 1,82 |
| | | Perception.Vision.Shape | 1,96 |
| | | Action.Observation | 2,15 |
| | | Cognition.Spatial | 2,24 |
| | | Perception.Vision.Motion | 2,59 |
| | | Action.Imagination | 2,82 |
| | | Action.Motor Learning | 5,3 |
| | FEF | Cognition.Spatial | 2,24 |
| | | Action.Execution | 2,52 |
| | | Action.Imagination | 2,89 |
| | | Perception.Vision.Motion | 3,44 |
| | 6d | Action.Imagination | 2,89 |
| | | Action.Execution | 4,27 |
| | | Action.Motor Learning | 6,43 |
| **Temporal** | STSva | Cognition.Memory.Explicit | 1,98 |
| | | Cognition.Language.Semantics | 2,36 |
| | | Cognition.Social Cognition | 4,32 |
| | | Emotion.Valence | 5,34 |
| | | Cognition.Language | 5,58 |
| | A5 | Cognition.Music | 1,87 |
| | | Cognition.Language | 2,78 |
| | | Action.Execution.Speech | 2,96 |
| | | Cognition.Language.Speech | 3,31 |
| | | Cognition.Language.Phonology | 3,76 |
| | | Perception.Audition | 4,26 |
| | | Action.Motor Learning | 4,47 |
| | STSdp | Cognition.Language.Semantics | 1,73 |
| | | Cognition.Language.Speech | 2,11 |
| | | Cognition.Social Cognition | 2,15 |
| | | Perception.Audition | 2,6 |
| | | Cognition.Language | 3,95 |
| | TE1m | Cognition.Language.Semantics | 2,31 |
| | TE1p | Cognition.Language.Speech | 2,34 |
| | | Cognition.Language.Orthography | 2,41 |
| | | Cognition.Language.Semantics | 2,48 |
| | | Cognition.Language.Phonology | 3,36 |
| | PHT | Cognition.Language.Semantics | 1,82 |
| | | Introception.Sexuality | 2,92 |
| | | Action.Observation | 4,46 |
| | TPOJ1 | Cognition.Language.Semantics | 1,94 |
| | | Cognition.Language.Speech | 2,06 |
| | | Perception.Audition | 2,53 |
| | | Emotion.Valence | 3,84 |
| | STV | Cognition.Social Cognition | 2,4 |
| | | Cognition.Music | 2,46 |
| | | Emotion.Valence | 3,96 |
| | TE1a | Cognition.Language.Semantics | 1,92 |
| | | Cognition.Language.Phonology | 2,86 |
| | | Cognition.Social Cognition | 4,63 |
| | | Cognition.Memory | 4,87 |
| | | Cognition.Language | 5,83 |
| | TE2a | No significant effects | |
| **Parietal** | PFt | Cognition.Language.Semantics | 2,39 |
| | | Cognition.Language.Phonology | 3 |
| | PGi | Cognition.Memory.Explicit | 1,7 |
| | | Cognition.Social Cognition | 4,48 |

**RIGHT HEMISPHERE**

| *lobe* | *area* | *domain* | *LR* |
|---|---|---|---|
| **Temporal** | TE1p | No significant effects | |
| | PHT | Emotion.Negative.Disgust | 3,67 |
| | TPOJ2 | Action.Observation | 3,74 |
| | | Emotion.Negative.Disgust | 4,34 |
| | TPOJ3 | Cognition.Social Cognitio | 3,16 |
| | | Perception.Vision.Shape | 3,67 |
| | | Cognition.Spatial | 4,62 |
| | TPOJ1 | Cognition.Social Cognitio | 2,61 |
| | | Perception.Audition | 2,61 |
| | TE1a | Cognition.Memory.Explici | 1,94 |
| | | Emotion.Negative | 2,35 |
| | | Cognition.Social Cognitio | 3,83 |
| | | Perception.Audition | 3,85 |
| | | Cognition.Language | 4,99 |
| | | Emotion.Valence | 5,98 |
| | STSdp | Cognition.Language.Sema | 1,94 |
| | | Cognition.Language.Spee | 2,57 |
| | | Emotion.Positive.Happine | 3,71 |
| | | Perception.Audition | 3,79 |
| | STV | Cognition.Social Cognitio | 3,34 |
| | PHT | Cognition.Spatial | 2,42 |
| | | Perception.Vision.Shape | 2,88 |
| | | Interoception.Secuality | 3,3 |
| | | Action.Observation | 4,44 |
| | FST | Perception.Vision | 2,41 |
| | | Emotion.Negative | 2,66 |
| | | Perception.Vision.Motion | 3,76 |
| | | Action.Observation | 3,94 |
| **Parietal** | PGs | Cognition.Social Cognitio | 2,6 |
| | PFm | Cognition.Reasoning | 1,51 |
| | | Perception.Somesthesis.F | 1,86 |
| | | Cognition.Memory.Workir | 1,87 |

**Supplementary table 1.** Functional decoding of all areas with minKL values>1.2 in the human:chimpanzee comparison. LR=likelihood ratio.

***HUMAN : MACAQUE***

**LEFT HEMISPHERE**

| *lobe* | *area* | *domain* | *LR* |
|---|---|---|---|
| **Frontal** | p47r | No significant effects | |
| | IFSa | Cognition.Language.Semantics | 2,33 |
| | | Cognition.Language.Orthography | 2,47 |
| | | Cognition.Language.Syntax | 3,01 |
| | | Cognition.Language.Phonology | 3,21 |
| | 45 | Cognition.Memory.Explicit | 1,63 |
| | | Emotion.Negative | 1,86 |
| | | Cognition.Language.Speech | 1,96 |
| | | Cognition.Language.Phonology | 2 |
| | | Cognition.Language.Semantics | 2,88 |
| | | Cognition.Language | 3,36 |
| | | Cognition.Language.Syntax | 4,36 |
| | 46 | Cognition.Reasoning | 1,55 |
| | | Cognition.Memory.Working | 1,67 |
| | | Cognition.Language.Speech | 1,67 |
| | | Cognition.Memory.Working | 2,04 |
| | | Cognition.Language.Semantics | 2,15 |
| | | Cognition.Language.Phonology | 2,47 |
| | p9-46v | Cognition.Language.Speech | 1,67 |
| | | Cognition.Memory.Working | 2,04 |
| | | Cogntiion.Language.Semantics | 2,15 |
| | | Cognition.Language.Phonology | 2,47 |
| | a9-46v | Cognition.Memory.Working | 2,16 |
| **Temporal** | STSva | Cognition.Memory.Explicit | 1,98 |
| | | Cognition.Language.Semantics | 2,36 |
| | | Cognition.Social Cognition | 4,32 |
| | | Emotion.Valence | 5,34 |
| | | Cognition.Language | 5,58 |
| | A5 | Cognition.Music | 1,87 |
| | | Cognition.Language | 2,78 |
| | | Action.Execution.Speech | 2,96 |
| | | Cognition.Language.Speech | 3,31 |
| | | Cognition.Language.Phonology | 3,76 |
| | | Perception.Audition | 4,26 |
| | | Action.Motor Learning | 4,47 |
| | STSdp | Cognition.Language.Semantics | 1,73 |
| | | Cognition.Language.Speech | 2,11 |
| | | Cognition.Social Cognition | 2,15 |
| | | Perception.Audition | 2,6 |
| | | Cognition.Language | 3,95 |
| | TE1m | Cognition.Language.Semantics | 2,31 |
| | TE1p | Cognition.Language.Speech | 2,34 |
| | | Cognition.Language.Orthography | 2,41 |
| | | Cognition.Language.Semantics | 2,48 |
| | | Cognition.Language.Phonology | 3,36 |
| | PHT | Cognition.Language.Semantics | 1,82 |
| | | Introception.Sexuality | 2,92 |
| | | Action.Observation | 4,46 |
| | TPOJ1 | Cognition.Language.Semantics | 1,94 |
| | | Cognition.Language.Speech | 2,06 |
| | | Perception.Audition | 2,53 |
| | | Emotion.Valence | 3,84 |
| | STV | Cognition.Social Cognition | 2,4 |
| | | Cognition.Music | 2,46 |
| | | Emotion.Valence | 3,96 |
| **Parietal** | PFm | Cognition.Reasoning | 1,5 |
| | | Cognition.Memory.Working | 1,59 |
| | | Cognition.Social Cognition | 2,05 |
| | PGs | Cognition.Memory.Explicit | 1,88 |
| | IP1 | Cognition.Reasoning | 1,81 |
| | MIP | Cognition.Memory.Working | 1,79 |
| | | Cognition.Language.Orthography | 2,3 |
| | | Perception.Vision.Shape | 2,45 |
| | | Cognition.Spatial | 2,91 |
| | LIPd | Perception.Vision | 1,87 |
| | | Cognition.Language.Orthography | 2,54 |
| | | Perception.Vision.Color | 3,55 |
| | VIP | Action.Execution | 2,54 |
| | | Perception.Vision.Motion | 3,96 |
| | 7Am | Perception.Vision.Motion | 2,44 |
| | | Cognition.Spatial | 2,7 |
| | | Action.Imagination | 3,2 |
| | 7Pl | Cognition.Attention | 1,64 |
| | | Cognition.Memory.Working | 2,04 |
| | | Perception.Vision.Shape | 2,63 |
| | | Cognition.Spatial | 2,98 |
| | | Perception.Vision.Color | 3,88 |
| | | Perception.Vision.Motion | 4,9 |
| | 7Pm | Perception.Vision.Motion | 3,23 |

**RIGHT HEMISPHERE**

| *lobe* | *area* | *domain* | *LR* |
|---|---|---|---|
| **Frontal** | 4 | Action.Execution | 3,17 |
| | | Action.Execution.Speech | 3,63 |
| | | Action.Motor Learning | 3,71 |
| | | Interoception | 3,73 |
| | 6mp | Action.Execution | 2,51 |
| | OFC | Emotion.Positive | 4,57 |
| | 10v | Cognition.Reasoning | 2,29 |
| | | Emotion.Positive.Reward/Gair | 3,17 |
| **Temporal** | TGd | Cognition.Memory.Explicit | 2,3 |
| | | Emotion.Positive | 2,94 |
| | | Emotion.Valence | 3,49 |
| | | Cognition.Social Cognition | 3,57 |
| | | Emotion.Negative.Sadness | 4,67 |
| | | Emotion.Negative.Anger | 4,79 |
| | | Emotion.Positive.Happiness | 4,9 |
| | | Emotion.Negative.Disgust | 5 |
| | Pir | Emotion.Negative | 2,48 |
| | | Emotion.Negative.Anger | 3,31 |
| | | Interoception.Sexuality | 3,34 |
| | | Emotion.Negative.Sadness | 4,43 |
| | | Emotion.Negative.Fear | 4,6 |
| | | Perception.Olfaction | 13,55 |
| | TE1p | No significant effects | |
| | PHT | Emotion.Negative.Disgust | 3,67 |
| | TPOJ2 | Action.Observation | 3,74 |
| | | Emotion.Negative.Disgust | 4,34 |
| | TPOJ3 | Cognition.Social Cognition | 3,16 |
| | | Perception.Vision.Shape | 3,67 |
| | | Cognition.Spatial | 4,62 |
| | TPOJ1 | Cognition.Social Cognition | 2,61 |
| | | Perception.Audition | 2,61 |
| **Parietal** | PGi | Cognition.Social Cognition | 4,13 |
| | PGs | Cognition.Social Cognition | 2,6 |
| | IP1 | Cognition.Reasoning | 1,99 |
| | | Cognition.Memory.Working | 2,41 |
| | PFm | Cognition.Reasoning | 1,51 |
| | | Perception.Somesthesis.Pain | 1,86 |
| | | Cognition.Memory.Working | 1,87 |
| | LIPd | Cognition.Attention | 1,73 |
| | | Cognition.Reasoning | 1,9 |
| | | Perception.Vision | 1,98 |
| | | Action.Inhibition | 2,58 |
| | | Cognition.Spatial | 3,08 |
| | | Action.Observation | 3,78 |
| | | Perception.Vision.Color | 4,82 |
| | LIPv | Cognition.Attention | 1,48 |
| | | Cognition.Memory.Working | 2,07 |
| | | Perception.Vision.Shape | 2,96 |
| | | Action.Observation | 3,38 |
| | | Perception.Vision.Motion | 4,01 |
| | | Cognition.Spatial | 4,44 |
| | 7PC | Action.Execution | 1,88 |
| | | Perception.Vision.Motion | 3,05 |
| | | Action.Motor Learning | 8,97 |
| | 7AL | Action.Execution | 2,12 |
| | | Action.Imagination | 4,49 |
| | | Cognition.Spatial | 4,71 |
| | | Action.Motor Learning | 16 |
| | 7Am | Cognition.Memory.Working | 2,59 |
| | | Action.Imagination | 3,5 |
| | | Perception.Vision.Motion | 3,68 |
| | 7Pm | Cognition.Memory.Working | 2,17 |
| | | Action.Inhibition | 2,77 |
| | | Perception.Vision.Motion | 3,06 |
| | PCV | No significant effects | |
| | 5L | Emotion.Positive.Happiness | 9,19 |
| | 5mv | No significant effects | |
| | 23c | Emotion.Negative | 3,21 |
| | | Emotion.Positive.Happiness | 4,66 |
| | 3b | Action.Execution | 3,06 |
| | | Perception.Gustation | 3,17 |
| | | Action.Execution.Speech | 4,99 |

**Supplementary table 2.** Functional decoding of all areas with minKL values>1.2 in the human:macaque comparison. LR=likelihood ratio.

### Supplementary material references


1. Van Essen, D. C. *et al.* The WU-Minn Human Connectome Project: an overview. *Neuroimage* **80**, 62–79 (2013).
2. Glasser, M. F. *et al.* The minimal preprocessing pipelines for the Human Connectome Project. *Neuroimage* **80**, 105–124 (2013).
3. Sotiropoulos, S. N. *et al.* Effects of image reconstruction on fiber orientation mapping from multichannel diffusion MRI: reducing the noise floor using SENSE. *Magn. Reson. Med.* **70**, 1682–1689 (2013).
4. Jenkinson, M., Beckmann, C. F., Behrens, T. E. J., Woolrich, M. W. & Smith, S. M. FSL. *Neuroimage* **62**, 782–790 (2012).
5. Chen, X. *et al.* Brain aging in humans, chimpanzees (Pan troglodytes), and rhesus macaques (Macaca mulatta): magnetic resonance imaging studies of macro- and microstructural changes. *Neurobiol. Aging* **34**, 2248–2260 (2013).
6. Autrey, M. M. *et al.* Age-related effects in the neocortical organization of chimpanzees: gray and white matter volume, cortical thickness, and gyrification. *Neuroimage* **101**, 59–67 (2014).
7. Bryant, K. L. *et al.* Organization of extrastriate and temporal cortex in chimpanzees compared to humans and macaques. *Cortex* **118**, 223–243 (2019).
8. Bryant, K. L., Li, L., Eichert, N. & Mars, R. B. A comprehensive atlas of white matter tracts in the chimpanzee. *PLoS Biol.* **18**, e3000971 (2020).
9. Andersson, J. L. R., Skare, S. & Ashburner, J. How to correct susceptibility distortions in spin-echo echo-planar images: application to diffusion tensor imaging. *Neuroimage* **20**, 870–888 (2003).
10. Behrens, T. E. J., Berg, H. J., Jbabdi, S., Rushworth, M. F. S. & Woolrich, M. W. Probabilistic diffusion tractography with multiple fibre orientations: What can we gain? *Neuroimage* **34**, 144–155 (2007).
11. Li, L. *et al.* Chimpanzee (Pan troglodytes) precentral corticospinal system asymmetry and handedness: a diffusion magnetic resonance imaging study. *PLoS One* **5**, e12886 (2010).
12. Donahue, C. J., Glasser, M. F., Preuss, T. M., Rilling, J. K. & Van Essen, D. C. Quantitative assessment of prefrontal cortex in humans relative to nonhuman primates. *Proc. Natl. Acad. Sci. U. S. A.* **115**, E5183–E5192 (2018).
13. Van Essen, D. C. Surface-based atlases of cerebellar cortex in the human, macaque, and mouse. *Ann. N. Y. Acad. Sci.* **978**, 468–479 (2002).
14. Mars, R. B., Passingham, R. E. & Jbabdi, S. Connectivity Fingerprints: From Areal Descriptions to Abstract Spaces. *Trends Cogn. Sci.* **22**, 1026–1037 (2018).
15. Warrington, S. *et al.* XTRACT - Standardised protocols for automated tractography in the human and macaque brain. *Neuroimage* **217**, 116923 (2020).
16. Fox, P. T. & Lancaster, J. L. Opinion: Mapping context and content: the BrainMap model. *Nat. Rev. Neurosci.* **3**, 319–321 (2002).
17. Glasser, M. F. *et al.* A multi-modal parcellation of human cerebral cortex. *Nature* **536**, 171–178 (2016).
18. Eickhoff, S. B. *et al.* Co-activation patterns distinguish cortical modules, their connectivity and functional differentiation. *Neuroimage* **57**, 938–949 (2011).
19. Genovese, C. R., Lazar, N. A. & Nichols, T. Thresholding of statistical maps in functional neuroimaging using the false discovery rate. *Neuroimage* **15**, 870–878 (2002).
20. Bailey, P., Bonin, G. V. & McCulloch, W. S. *The Isocortex of the Chimpanzee.* (Univ. of Illinois Press, Oxford, England, 1950).
21. Percheron, G., François, C. & Pouget, P. What makes a frontal area of primate brain the frontal eye field? *Front. Integr. Neurosci.* **9**, 33 (2015).
22. Petrides, M. Lateral prefrontal cortex: architectonic and functional organization. *Philos. Trans. R. Soc. Lond. B Biol. Sci.* **360**, 781–795 (2005).

23. Neubert, F.-X., Mars, R. B., Thomas, A. G., Sallet, J. & Rushworth, M. F. S. Comparison of human ventral frontal cortex areas for cognitive control and language with areas in monkey frontal cortex. *Neuron* **81**, 700–713 (2014).
24. Pandya, D. N. & Seltzer, B. Intrinsic connections and architectonics of posterior parietal cortex in the rhesus monkey. *J. Comp. Neurol.* **204**, 196–210 (1982).
25. Mars, R. B. *et al.* On the relationship between the 'default mode network' and the 'social brain'. *Front. Hum. Neurosci.* **6**, (2012).
26. Schurz, M., Tholen, M. G., Perner, J., Mars, R. B. & Sallet, J. Specifying the brain anatomy underlying temporo-parietal junction activations for theory of mind: A review using probabilistic atlases from different imaging modalities. *Hum. Brain Mapp.* **38**, 4788–4805 (2017).
27. Milham, M. P. *et al.* An Open Resource for Non-human Primate Imaging. *Neuron* **100**, 61–74.e2 (2018).
